# Links Between Gut Microbiota of Uruguayan Infants, Breastmilk Composition, and Maternal Factors During Exclusive Breastfeeding

**DOI:** 10.64898/2025.12.16.691825

**Authors:** Laura Herrera-Astorga, Cecilia Mathó, Joaquín Pereira-Pagola, Julieta Pomi, Lucía Bilbao, Carolina Farías, Arturo Puyol, Claudio Rodríguez-Camejo, José Roberto Sotelo-Silveira, Ana Hernández

**Author notes:** These authors contributed equally (Laura Herrera-Astorga, Cecilia Mathó, Joaquín Pereira-Pagola).

## Abstract

**Background:** Given the links between early gut microbiota, breastfeeding, and maternal physiology, we characterized the intestinal microbiota of infants during exclusive breastfeeding (mean=5.4 months), examining its associations with breastmilk composition, fecal IgA, and maternal factors in Uruguayan primiparous mothers, with vaginal (V, n=20) or cesarean-section delivery (CS, n=14).

**Methods:** We analyzed fecal microbiota via 16S rRNA V4 sequencing and quantified stool IgA, breastmilk components (antibodies, hormones, macronutrients) using ELISA and standard biochemical methods.

**Results:** Principal Coordinates Analysis showed separation by delivery mode (PCo1:44.6%, PCo2:14.2%). Samples of CS-group displayed lower relative abundance (RA %) of Bacteroidetes/Firmicutes ratio, higher RA of Proteobacteria (50.7%vs35.5%), and decreased Bacteroides (2.5%vs31.8%) than V-group. Biochemical parameters didn’t differ between groups. In the V-group, milk and fecal IgA correlated (r=+0.47), as did Bifidobacterium RA with milk IgA (r=+0.47).

Fat content was associated with different microbial taxa in both groups. Only in CS-group milk carbohydrates correlated with Bifidobacterium (r=-0.679) and maternal stress with Flavonifractor in CS-group (r=+0.461).

**Conclusion:** Results indicate delivery mode can exert persistent impact on infant gut microbiota until the introduction of complementary feeding. Differences in correlation patterns between groups suggest distinct regulatory mechanisms of microbiota, possibly linked to physiological processes that differ according to delivery mode.

**Impact:**

- This is the first study to assess the gut microbiota composition of exclusively breastfed infants born to Uruguayan mothers, in parallel with analyses of breast milk composition and maternal perceived stress
- The mode of delivery was associated with differences in gut microbiota composition at a mean age of 5.4 months.
- Different associations between milk composition and maternal perceived stress with predominant microbial taxa were found according to the type of delivery.
- These findings provide a basis for studies on microbiota regulatory mechanisms influenced by maternal physiology according to delivery mode

## INTRODUCTION

The gut microbiota is central to human health, and its establishment in early life is critical for the health outcome during infancy and beyond ^1,2^. Early microbiota composition has a strong relationship with the development of adult microbiota and directly influences metabolism and immune development^3^. The early diversity and colonization patterns of gut microbiota have been associated with the onset of immune-related pathologies during infancy ^4,5^.

Several intrinsic and environmental perinatal factors contribute to the composition of early gut microbiota, with mode of birth and feeding pattern being major determinants^6–10^. Cesarean section delivery (CS) has been shown to disrupt the natural transmission of maternal vaginal and gut microbiota to offspring ^11,12^. As a result, infants delivered by CS may miss early exposure to microbial species that Yu contribute to immune maturation and gut development ^13,14^. The impact of CS is further influenced by the use of intrapartum antibiotics, and whether the CS was planned or if it occurred after onset of active labor ^6,8,15^.

These alterations in early microbial colonization may have long-term health outcomes. Infants born by CS are at increased risk of metabolic diseases and infections during the first months of life, with microbiota composition shift proposed as a contributing factor to these risks. ^6,16^

However, microbial transmission beyond the delivery mode also strongly contributes to establishing the infant gut microbiota^17^. Breastmilk provides a complex community of microorganisms that will colonize the infant gut and supplies several components that influence the microbiota profile ^9,18,19^. Several milk prebiotics can favor growth of bacteria that produce beneficial metabolites. Nonetheless, the impact of breastmilk composition on the early microbiota profile has been mainly studied in relation with the profile of human milk oligosaccharides, the main prebiotic components in breastmilk^20,21^. Homeostasis of gut bacteria is also regulated by the interaction with secretory antibodies (IgA and IgM), which are only provided by breastmilk in the initial postnatal stages^22^.

However, breastmilk composition is highly variable among women of different geographic regions, with the supply of some components being dependent on genetics and environmental factors^23–25^. This highlights the need to further investigate diverse populations to capture variations in the early-life factors shaping gut microbiota. Despite this, many regions remain understudied, leaving a critical gap that must be addressed. Here, we present the first study of gut microbiota in healthy Uruguayan infants at 5.5 months of age, who have been exclusively breastfed since delivery. We analyzed the association of the relative abundance (RA) of taxa with: i. maternal factors, *ii*. the level of milk macronutrients and functional components that are known to impact microbiota composition (secretory antibodies, cortisol, adiponectin and leptin); iii. the level of IgA in the infant’s stool.

## MATERIALS AND METHODS

### Study design and participants

This study involves a group of Uruguayan mother-infant dyads who maintained exclusive breastfeeding until the introduction of complementary feeding (between 5-6 months). The volunteers were contacted by researchers between May 2020 to April 2022, and all were living in the metropolitan area of Montevideo. The recruitment strategies included direct communication with breastfeeding support organizations and institutional social networks to provide information about our aims. The researchers contacted the mothers interested in participating and explained in detail the logistics of providing biological samples and filling in the questionnaires.

The inclusion criteria were defined as: primiparous women with no significant morbidities, mothers of healthy term infants, being breastfeed from the first hours after parturition without formula administration after discharge. The following exclusion criteria were considered: i. multiple pregnancy; ii. preterm or low birth weight; iii. maternal chronic or infectious diseases; iv. smoking; v. child health complications; vi. gynecological-obstetric complications.

At the time of enrollment, the mothers were asked about the higher education level, age at parturition, gestational age, type of delivery, following a restrictive diet, pre-pregnancy body mass index (BMI: weight/height, Kg/m^2^) and weight gain during pregnancy. In the case of mothers who had a cesarean section, they were asked whether it was scheduled and whether they received antibiotics during the procedure.

Infant core anthropometric measures (body weight and length) were obtained from medical records starting at delivery and during the first year. The Spanish Gastroenterology Hepatology and Nutrition Society application was used to calculate the percentiles (P) for weight for length ratio (W/L) at 6- and 12-month age according to WHO 2006-2007 recommendations^26^.

### Perceived stress evaluation

At the time of sample collection, the mothers filled in a standard questionnaire to evaluate the stress perceived over the past month. We used the validated Perceived Stress Scale (PSS14) based on 14 questions ^27,28^. Briefly, the items adopt a five-option response format labeled as follows: never, almost never, sometimes, fairly often, and very often. To obtain the score, integer values are assigned to each response option as follows: 0 = never; 1 = almost never; 2 = sometimes; 3 = often; and 4 = very often. The total score is calculated by summing the value assigned to the response chosen for each item. Higher scores indicate greater perceived stress.

#### Milk samples

Mothers were asked to donate a milk sample (2-10mL) between 5 and 6 months postpartum during exclusive breastfeeding, just before the introduction of complementary foods. Hindmilk samples were obtained after feeding their infants with a breastmilk pump in the morning (between 7:00 and 10:00 AM). The milk was collected in sterile polypropylene containers provided by researchers. Samples were immediately frozen at −20°C in home freezers until collected by the researchers. Once in the laboratory, the samples were conserved at −80°C until analysis. For some determinations, the aqueous fraction of milk (AF) was separated by two sequential centrifugation steps (1.000 rpm for 10 min at 4°C and 10.000 rpm for 30 min at 4°C) to remove cells, debris and fat^29^. The AF was aliquoted and stored at −80°C until use to measure milk components as indicated below.

#### Stool samples

In parallel to the milk sample collection, the mothers were instructed to collect a sample of the infant’s stool. The researchers provided the sterile container and swab to collect the sample. The specimens collected by mothers were immediately stored at −20 °C in home freezers and retrieved by the researchers within a few days. Once in the laboratory, the samples were conserved at −80 °C until processing.

### Macronutrients in breastmilk

Protein concentration in the AF was determined with the Micro BCA™ Protein Assay Kit (Thermo Fisher Scientific, Waltham, MA) following the manufacturer’s instructions for microplates. Standard curves were constructed using the AF of mature milk with protein concentration evaluated by the Kjeldahl method (11.4 mg/mL)^30^.The carbohydrate content of the AF was estimated using the 3,5-dinitrosalicylic acid method and D-lactose was used as a standard ^31^.The crematocrit (defined as the percentage of cream of the unfractionated human milk, %C) which correlates with fat content and energy, was measured in triplicate samples of whole milk^32^.

### Hormones in breastmilk

Cortisol concentrations were determined via Cortisol Saliva Luminescence Immunoassay Kit (IBL-International, Hamburg, Germany). This assay was previously adapted to the analysis of milk samples^33^. Briefly, 500 μL of chilled dichloromethane was added to 100 µL of whole milk samples. After vortexing, the samples were maintained in ice for 10 min, and then centrifuged at 1500 g for 5 min. After removing the AF, the solution was evaporated until dry in a fume hood at room temperature. The content was resuspended with 50 μL of distilled water and the samples were analyzed in duplicate according to the kit instructions. Relative luminescence units were recorded within 10 min by using a Fluostar Optima Reader (BMG, Ortenberg, Germany).

The leptin and adiponectin concentrations in the AF of milk samples were measured by using the respective enzyme-linked immunosorbent assay (ELISA) kits: Human Leptin DuoSet ELISA and Human Adiponectin/Acrp30 DuoSet ELISA (R&D Systems, Minneapolis, MI) according to the manufacturer’s instructions. For leptin assay, the samples were diluted x3 in dilution buffer and undiluted samples were used for adiponectin and ghrelin analysis.

### Antibodies in breastmilk

Total immunoglobulins IgA, IgM, and IgG in milk were quantified by customized capture enzyme-linked immunosorbent assay (ELISA), as previously described^29,30^. Briefly, microplates (Greiner Bio-One, Frickenhause, Germany) were coated with the corresponding capture antibody in PBS overnight (4°C). Following blocking with 1% gelatin in phosphate-buffered saline (PBS) (1h, 37°C), milk samples AF or standard were added in 0.2% gelatin and 0.05% Tween-20 in PBS and incubated overnight (4°C). Then, samples were incubated for 1 h (37°C) with horseradish peroxidase (HRP)-conjugated antibodies. The color reaction was then developed using a solution of sodium acetate (pH:5.5) containing 1.3 mM H_2_O_2_ and 0.4 mM 3,3′,5,5′-tetramethylbenzidine. The reaction was stopped with 2N sulfuric acid and the optical density at 450 nm (OD_450_) was then registered. The capture antibodies used were rabbit anti-human-Ig Fc antibody (Sigma-Aldrich, St. Louis, MO) for IgA and IgM, and goat anti-human-Fcγ antibody (Abcam, Cambridge, UK) for IgG. For detection of IgA and IgM goat anti-human-Fcα-HRP (Sigma-Aldrich) and goat anti-human-Fcμ-HRP (Sigma-Aldrich) were used, respectively. Goat anti-human-Fcγ-HRP (Dako, Glostrup, Denmark) absorbed with goat serum was used for IgG detection. Standard curves were constructed with a commercial N Protein Standard SL (Siemens Healthcare Diagnostics Products GmbH, Marburg, Germany).

#### IgA in stool

For analysis of stool IgA, 1 g of each specimen was diluted with PBS (w/v) and an extract was obtained as previously described^34^. Briefly, after vortexing for 30 min at 4°C. the suspension was centrifuged at 13.000 rpm for 15 min. The supernatant was collected, diluted 1:1 and 1:5 with sample buffer and analyzed by ELISA as described above.

#### Stool DNA extraction

Total DNA was extracted from the babies’ stool samples with QIAamp Fast DNA Stool Mini Kit (#51604,Qiagen, Venlo, Netherlands), following the manufacturer’s instructions.

### 16S Library preparation from Stool DNA

The V4 hypervariable region of the 16S rRNA gene was amplified from DNA extracted from stool samples using the forward primer 520F (AYTGGGYDTAAAGNG) and reverse primer 802R (TACNNGGGTATCTAATCC)^35^, each incorporating sample-specific barcodes and sequencing adaptors to construct libraries, as previously described^36^. Libraries were quantified using a Qubit™ Fluorometer with the Qubit™ 1X dsDNA High Sensitivity Assay Kit (#Q33230, Thermo Fisher Scientific), and their fragment size distribution was assessed using a Bioanalyzer 2100 (Agilent Technologies, Santa Clara, USA) with the High Sensitivity DNA Kit.

### NGS sequencing

Samples were diluted to a final concentration of 50 pM and pooled in an equimolar ratio for template preparation. Template preparation and Ion 530™ Chip loading were carried out on the Ion Chef™ Instrument according to the manufacturer’s instructions. Sequencing was performed using the Ion GeneStudio™ S5 System with 850 flows, following the standard protocol. All sequencing procedures were conducted at the Plataforma de Secuenciación Masiva (NGS sequencing facility) of the Instituto de Investigaciones Biológicas Clemente Estable.

### Bioinformatic Analysis

For the automated analysis of data generated through high-throughput sequencing, a bioinformatics workflow was designed using Snakemake (v7.32.3) ^37^. This workflow management system enables the integration and execution of multiple tools in reproducible and scalable environments. The main goal was to develop an automated workflow for processing data derived from 16S rRNA metagenomic sequencing.

The workflow begins with module dedicated to filtering and quality control of raw reads, removing bases with errors and artifacts generated during sequencing. For this, the programs

FastQC^38^ and MultiQC^39^ are used, which create an initial report on the status and integrity of the sequences. Subsequently, quality filtering and artifact removal are performed using QIIME2^40^ plugins (quality-filter^41^ and VSEARCH^42^). In addition, the DADA2^43^ plugin is used to group the sequences into ASVs (Amplicon Sequence Variants).

Once the sequences are grouped into ASVs, taxonomy is assigned using QIIME 2’s feature-classifier plugin, with a pre-trained human stool-weighted classifier based on the SILVA database for 16S rRNA sequences ^44–46^. The annotated sequences are then exported in FASTA and CSV table formats.

From the taxonomically assigned sequences, the next module infers a phylogeny that is used to study the microbial diversity present in the samples. Alpha diversity analysis includes the calculation of richness-based metrics ^47^, evenness (Pielou’s evenness or pielou_e), and diversity (Faith Phylogenetic Diversity ^48^) along with rarefaction analysis. On the other hand, beta diversity is assessed using weighted UniFrac ^49^ metric, visualized through Principal Coordinates Analysis (PCoA). Group comparisons are also considered through analysis of microbial composition with bias correction ^50^.

The workflow was validated using previously obtained laboratory data, including controlled microbial communities and environmental samples, which allowed fine-tuning of analysis parameters and optimization of the system’s sensitivity and specificity. Positive and negative controls were also incorporated, along with comparison to host genome sequences to prevent false positives.

### Statistical analysis

We tested the normal distribution of experimental data of milk and stool composition using the Kolmogorov-Smirnov test. Since most variables did not follow a normal distribution, we used non-parametric statistics for analysis. Median and interquartile ranges (IQR: 25th and 75th percentiles) were used to present the results. Mann–Whitney U test was used for comparisons between groups. Spearman’s test was used to analyze the correlation between each taxon (with RA (%) ≥1 and prevalence >40%) and maternal–infant dyad variables (maternal age, BMI, weight gain, perceived stress, IgA in stool), as well as compositional data of breastmilk.

The statistical significance level was defined as 95% confidence and is indicated in the figures with asterisks: *p ≤0.05, **p ≤0.01, and ***p ≤0.001. GraphPad Prism 8.4.3 (demo) was used for statistical analysis and data representation.

We tested several multiple linear regression models with RA of genera showing a RA ≥1%. The outcome variables were selected according to the results of simple analysis by Spearman test (r>|0.2|). The type of birth was the cofactor in all models.

To assess differences in overall microbial community structure between infants born by vaginal delivery (V) and cesarean section (CS), a permutational multivariate analysis of variance (PERMANOVA) was performed on a weighted UniFrac distance matrix between samples^51^. PERMANOVA was used to test for differences in community composition between delivery modes, while permutational analysis of multivariate dispersions (PERMDISP) was applied to evaluate whether observed differences were driven by heterogeneity in within-group dispersion^52^.

## RESULTS

### Characteristics of the study group

A total of 34 mother–infant dyads were included in the study. All mothers were primiparous, lived with their partners, and 88% reported having tertiary education.

All infants were born at term (>37 weeks); twenty were delivered vaginally and fourteen by cesarean section, with half of the cesarean deliveries occurring after labor onset.

Exclusive breastfeeding was maintained for an average of 5.4 ± 0.4 months (SD), independently of the mode of delivery. No differences were found between the anthropometric data collected at the time of sample collection and at 1 year of age. The characteristics of infants and mothers are detailed in **Table 1**.

**Table 1.**
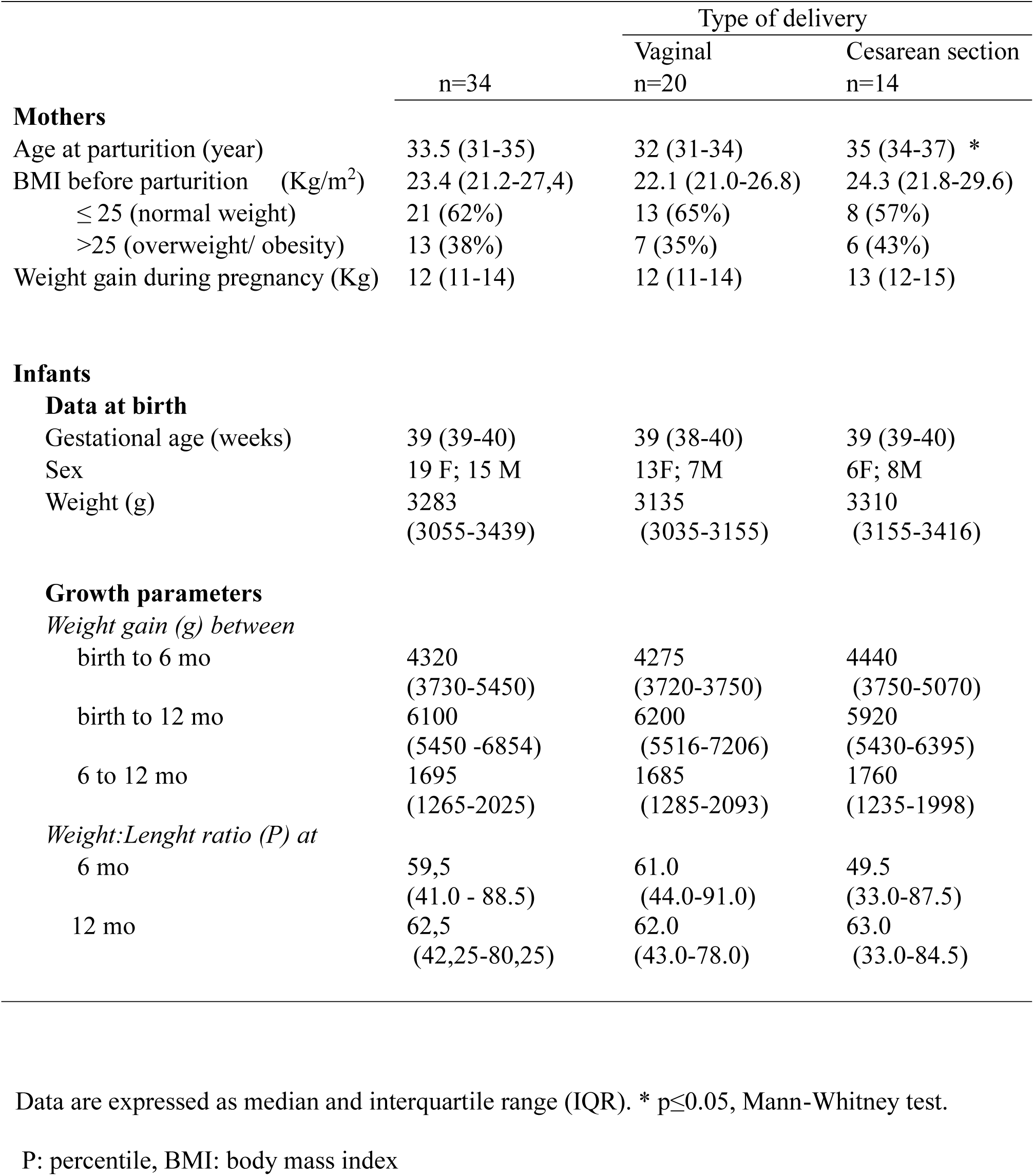
Characteristics of mothers and their infants of the study group.

### Infant gut microbiota composition

NGS sequencing of the 16S V4 region showed that the sequencing depth was adequate for all samples, as indicated by the rarefaction curves, which reached a clear plateau (**Supplementary Figure 1**).

Principal coordinates analysis (PCoA) based on the weighted UniFrac distance was performed to evaluate factors differentially affecting gut microbiota composition in our cohort. This analysis revealed that PCo1 and PCo2 explained 44.6% and 14.2% of the variance, respectively, and contributed to the separation of infants born by vaginal (V) or cesarean section (CS) delivery **(Figure 1)**.

**Figure 1.**
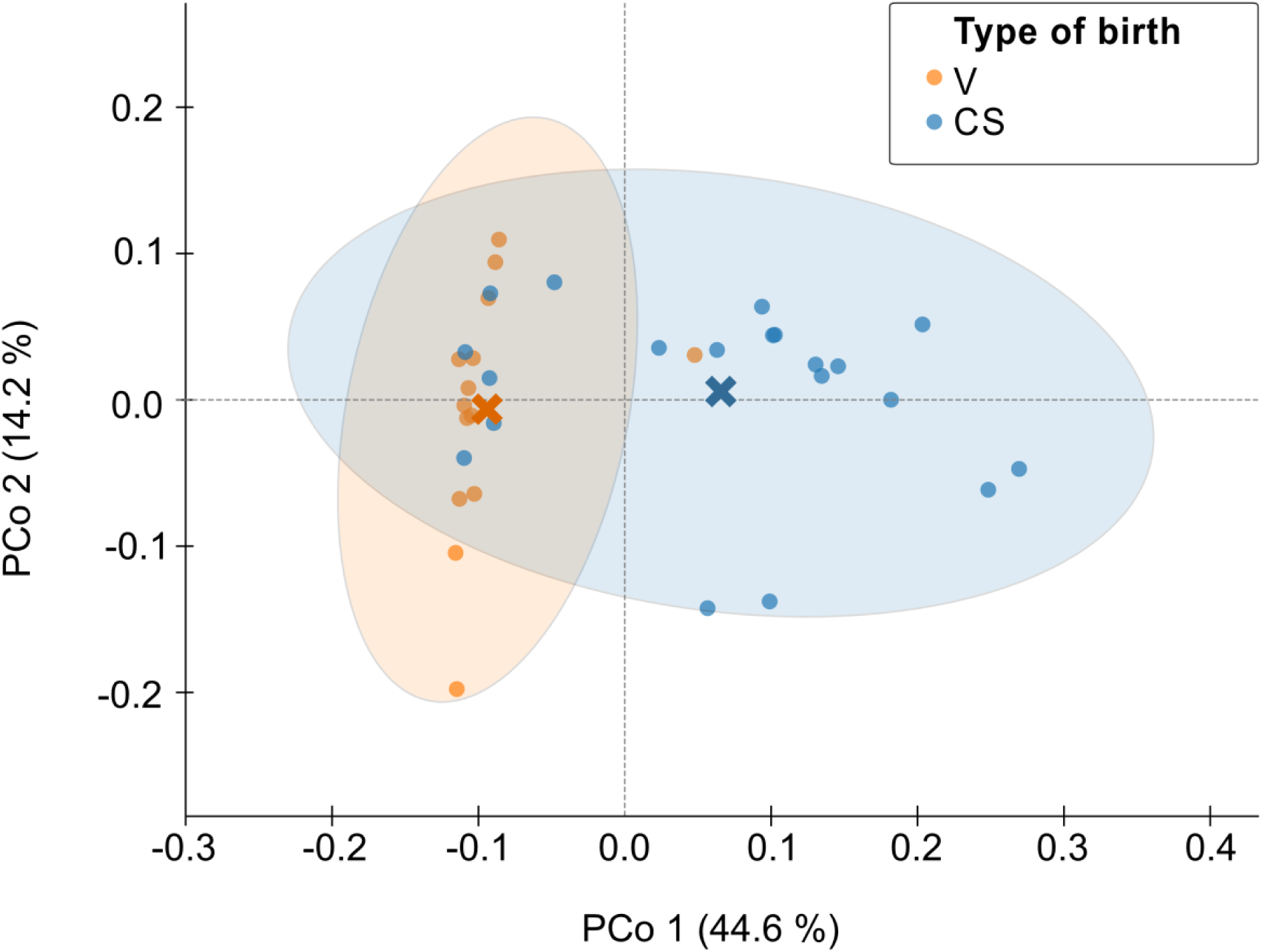
Principal coordinate analysis (PCoA) of infant stool microbiota. The PCoA was performed on a distance matrix derived from microbial abundance profiles for each sample (n=34), using weighted UniFrac as the distance metric. The first two principal coordinates (PCo1 and PCo2) account for 44.6% and 14.2% of the variability, respectively. Each dot represents an individual sample, and its color represents the delivery mode. Mean PCo values for each group along axes are indicated by crosses, and the ellipses represent 95% confidence intervals. PERMANOVA analysis confirmed a significant effect of delivery mode on microbial community structure (F = 8.47, p = 0.003, 300 permutations).

PERMANOVA analysis with 300 permutations confirmed a significant effect of delivery mode on microbial community structure (F = 8.47, p = 0.003). Although dispersion analysis (PERMDISP) indicated a moderate difference in variability between groups (F = 4.93, p = 0.073), the strong effect detected by PERMANOVA supports delivery mode as a major factor modulating beta diversity.

Given these results, subsequent analyses were performed considering two groups according to type of delivery. At the phylum level, Proteobacteria, Bacteroidetes, Firmicutes, and Actinobacteria were present in almost all samples, except that Bacteroidetes were missing in three CS infant samples (**Table 2**).

**Table 2.** Prevalence and relative abundances, RA of the dominant bacteria in infant stool.

|  | Prevalence (%) |  | RA (%) |  |  |
| --- | --- | --- | --- | --- | --- |
|  | V | CS | V | CS |  |
| **p_Bacteroidetes** | 100 | 79 | **38.5±26.6** | **2.6±9.4** | **** |
| f_Bacteroidaceae | 95 | 50 | 31.8±26.7 | 2.5±9.1 | **** |
| g_Bacteroides | 95 | 50 | 31.8±26.5 | 2.5±9.4 | **** |
| f_Tannerellaceae | 35 | 0 | 1.1±2.2 | - |  |
| g_Parabacteroides | 35 | 0 | 1.1±2.2 | - |  |
| **p_Firmicutes** | **100** | **100** | **20.6±17.3** | **38.5±22.9** | * |
| f_Clostridiaceae | 100 | 100 | 6.9±12.8 | 16.4±22.0 | * |
| g_Clostridium_sensu_stricto_1 | 100 | 100 | 6.9±13.1 | 16.4±22.4 |  |
| g_Flavonifractor | 60 | 57 | 1.5±4.1 | 1.5±2.5 |  |
| f_Oscillospiraceae | 60 | 57 | 1.5±4.0 | 1.5±2.5 |  |
| g_[Ruminococcus]_gnavus_group | 90 | 86 | 2.0±3.1 | 6.7±12.9 |  |
| f_Veillonellaceae | 100 | 100 | 4.8±7.8 | 9.9±13.52 |  |
| g_Veillonella | 100 | 100 | 4.6±7.1 | 9.7±13.5 |  |
| f_Lactobacillaceae | 80 | 0 | 3.2±9.5 | - |  |
| g_Lactobacillus | 40 | 0 | 3.0±9.4 | - |  |
| f_Lachnospiraceae | 100 | 93 | 2.5±3.3 | 8.7±14.0 |  |
| **p_Proteobacteria** | **100** | **100** | **35.5±18.7** | **50.7±20.1** | * |
| f_Enterobacteriaceae | 100 | 100 | 31.0±0.8 | 49.5±24.3 | * |
| g_Escherichia-Shigella | 100 | 100 | 26.1±20.0 | 37.1±28.3 |  |
| f_Morganellaceae | 35 | 0 | 2.2±6.5 | - |  |
| g_Morganella | 30 | 0 | 2.2±6.6 | - |  |
| f_Sutterellaceae | 40 | 0 | 1.6±6.1 | - |  |
| g_Sutterella | 40 | 0 | 1.6±6.2 | - |  |
| f_Pasteuralleceae | 0 | 13 | - | 1.1±2.5 |  |
| g_Conservatibacter | 0 | 86 | - | 1.0±2.5 |  |
| **p_Actinobacteria** | **100** | **100** | **7.0±7.7** | **4.9±7.9** |  |
| f_Bifidobacteriaceae | 95 | 100 | 6.7±7.6 | 4.7±7.7 |  |
| g_Bifidobacterium | 95 | 100 | 6.7±7.6 | 4.7±7.7 |  |

Data shows the prevalence and relative abundances, RA of the dominant bacteria at the phylum, family, or genus levels, detected in stool samples grouped according to the type of delivery: vaginal (V) or cesarean section (CS). Data of RA are expressed as mean (standard deviation). The asterisks indicate significant differences (Mann-Whitney test).

Comparative analysis of stool samples from V and CS infants showed significant differences in the RA of certain taxa at the phylum, family, and genus levels. As shown in **Figure 2a**, the phylum Bacteroidetes was found in significantly higher RA in the group of V infants. This difference was maintained in the Bacteroidaceae family and *Bacteroides* genus (**Figure 2b-c).** Moreover, within the phylum Bacteroidota, f_Tannarellaceae and *g_Parabactoides* were only detected in the stool samples of the V group, with an average prevalence of 35% **(Table 2).**

**Figure 2.**
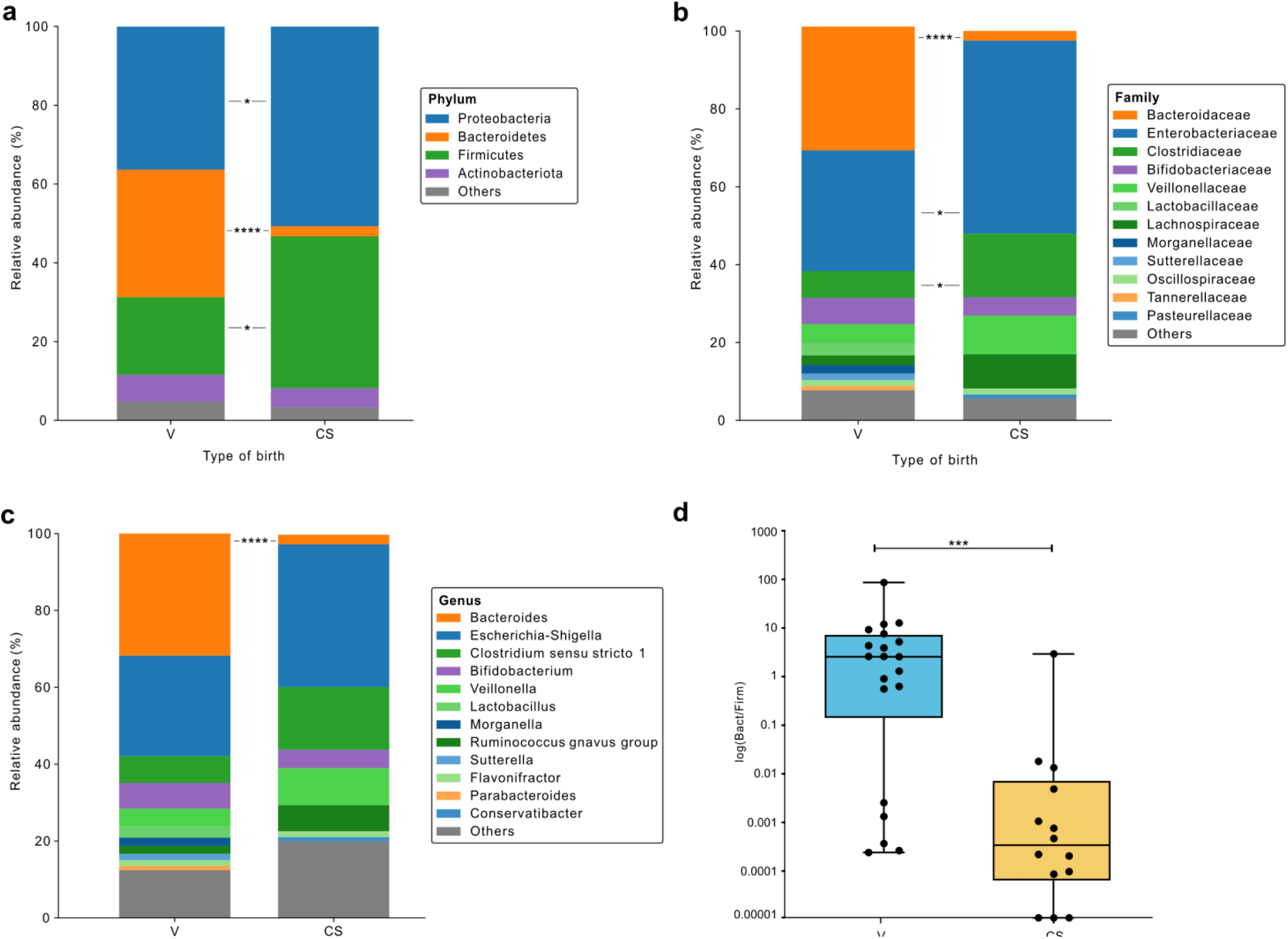
Comparison of the mean relative abundance (RA) of gut microbiota at the phylum (a), family (b) and genus (c) levels between the V and CS groups. Taxa with RA ≥1% are shown in the bar plot. Bacteria with RA below 1% are grouped as “others” and shown in grey color. The graph in panel d compares the ratio RA of Bacteroidota/Firmicutes phyla in samples corresponding to V and CS groups. The asterisks show significant differences (Mann-Whitney test).

Consistent with the mode of delivery, f_Lactobacillaceae and *g_Lactobacillus* were detected in 80% and 40% of the samples from the V infant group, respectively, but were not detected in children born by CS, either with or without previous labor.

The RA of the Proteobacteria phylum and Enterobacteria family were higher in the CS group. f_Enterobacteriaceae and g_*Escherichia-Shigella* were dominant taxa detected in all samples of both groups. In contrast, the families/genera Morganellaceae*/Morganella* and Sutterellaceae*/Sutterella* were only detected in samples of V infants, with prevalences ranging between 30 and 40%. Only the genus *Conservatibacter* was exclusively detected in 13% of the samples of the CS group **(Table 2).**

The ratio of RA of Bacteroidetes and Firmicutes was significantly different between groups, median: 2.9 vs 0.0003 for V and CS groups, respectively (**Figure 2d).**

The comparison of alpha indexes of diversity for phylogeny (Pielou) and evenness (Faith) showed no differences between groups V and CS **(Figure 3).**

**Figure 3.**
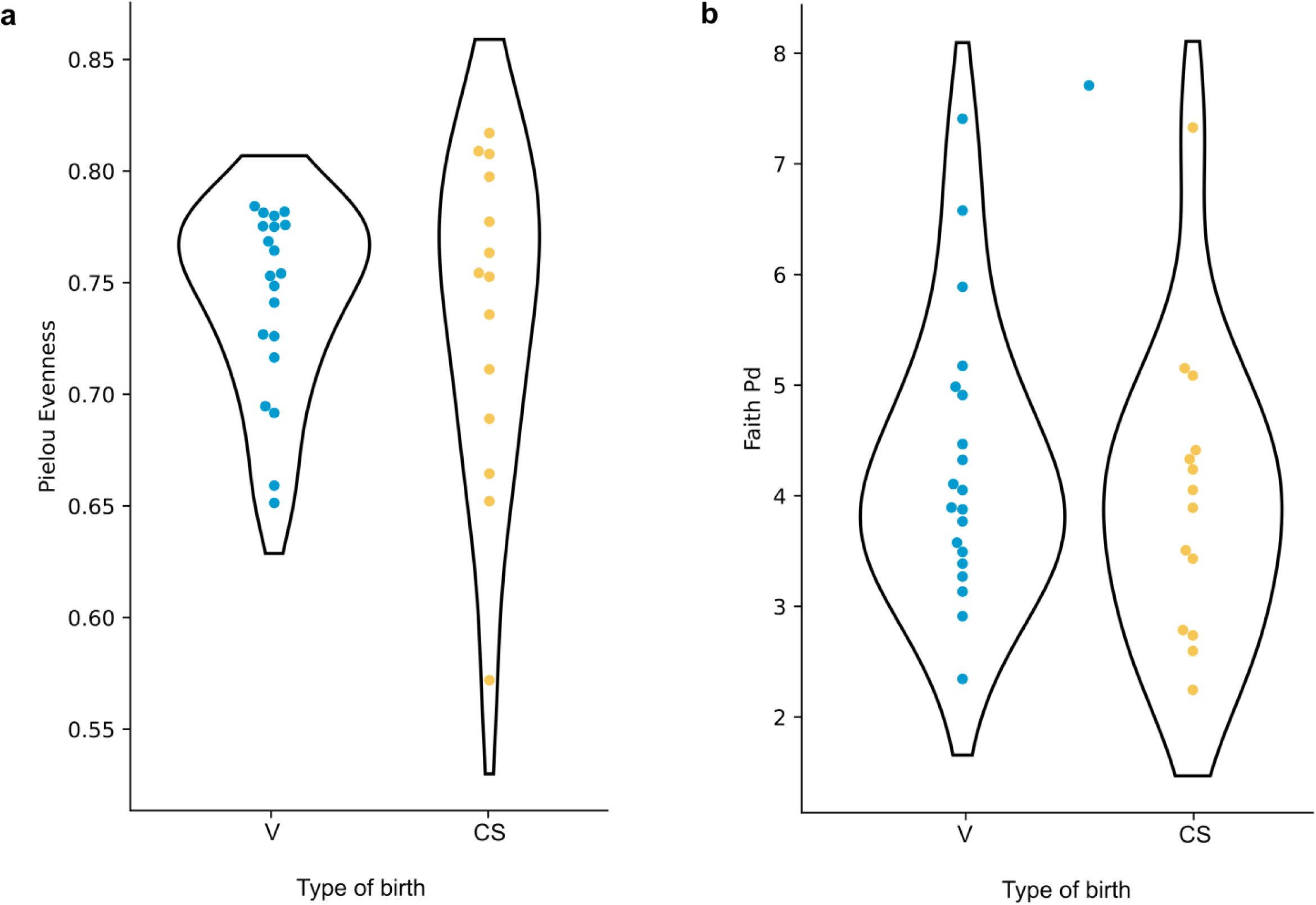
Violin plots show distributions of alpha-diversity by type of delivery for evenness (a) and phylogenetic (b) indices.

### Analysis of biochemical data and maternal perceived stress in relation to the mode of delivery

The content of macronutrients, antibodies and macronutrients in milk was similar between samples provided by mothers with V or CS delivery **(Figure 4 a-c).** Likewise, fecal IgA levels did not differ according to the mode of delivery **(Figure 4d).** Nevertheless, a significant correlation between the level of IgA in milk and stool, was only observed in the V group (r=+0.47, p=0.04).

**Figure 4.**
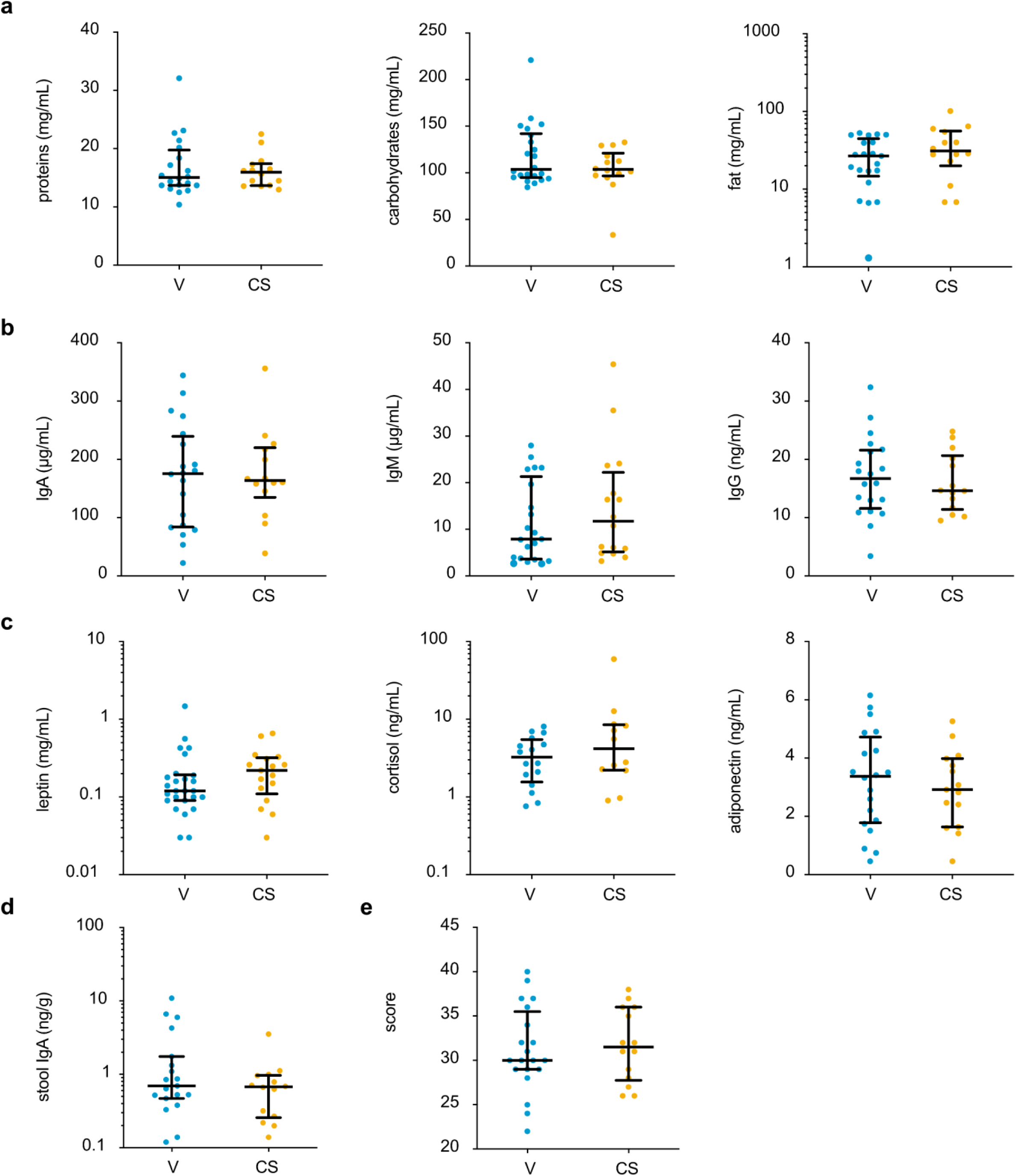
Comparison of experimental data according to the type of delivery. The graphs show the results for each individual sample: levels of macronutrients (a), antibodies (b) and hormones (c) in milk. Panel d shows IgA levels in the infant stool. The scores for the mother’s perceived stress according to PSS14 questionnaire completed at the time of sample collection are shown in panel e. Median and IQR values are indicated with bars. V: vaginal delivery, CS: cesarean section.

Regarding the perceived stress of mothers who had a vaginal delivery or a cesarean section, no significant differences were observed in the respective median scores **(Figure 4 e)**.

### Correlation analysis of microbiota composition in relation to mother–infant dyad parameters

Given the differences in RA profiles between the V and CS groups, we conducted a group-specific simple correlation analysis between microbiota composition (phylum, family, and genus RA), experimental variables, and maternal characteristics. We restricted the analysis to those taxa with RA ≥ 1% and present in ≥ 40% of the participants. The analysis of correlation with milk components showed significant correlations between milk fat and carbohydrate contents with different taxa, depending on the type of delivery **(Figure 5a)**. Regarding the content of hormones and microbiota composition, significant correlations were only observed with the leptin level in the CS group. In relation to the content of antibodies in milk, a significant positive correlation between IgA and RA of *g_Bifidobacterium* was observed in the V group. Additionally, a non-significant trend was observed between the RA of these taxa with stool IgA in this group (p=0.07) **(Figure 5b).**

**Figure 5.**
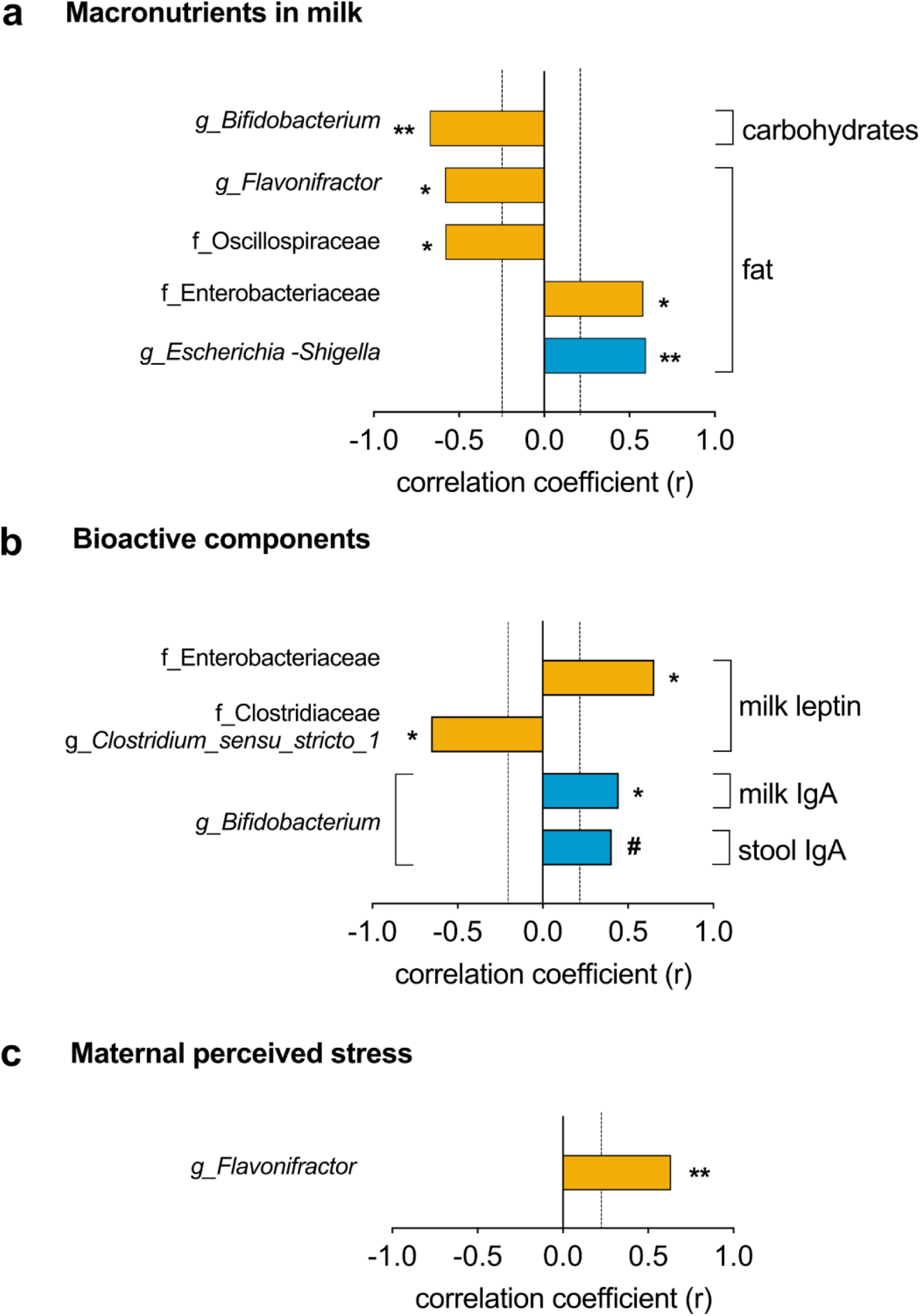
Spearman correlation analysis between the RA (%) of taxa (family and/or genus level) and experimental data of biochemical analysis of samples (a-b), and the score of perceived maternal stress (c) are depicted in the graphs. Taxa with relative abundance ≥1% and detected in at least 40% of samples were used for analysis. Significant correlations are indicated with asterisks, and a non-significant trend is indicated by #. The orange bars show data of the CS-group while the blue bars correspond to V-group.

Regarding the maternal perceived stress, in the CS group, we found a significant positive correlation with the RA of *g_Flavonifractor* **(Figure 5c)**.

No significant correlations were observed between the microbiota composition and maternal age, nor the parameters related to their body weight.

In parallel, we performed a multivariate analysis using data from the whole cohort (n=34), with microbiota genera as independent variables and birth type as a cofactor. To select the variables for the models, we previously performed a simple Spearman correlation analysis and considered those with r>|0.3| **(Supplementary Table 1).**

The significant correlations detected in simple correlation analyses within the V and CS groups were no longer observed when multivariable regression models were applied, after adjusting for birth type. The results obtained with the multiple analyses are detailed in **Table 3**.

**Table 3.** Multiple linear regressions modeling the association between microbiota and variables of mother age and milk components.

| Outcome variables | F | p | adjusted R-square | type of delivery | Beta coefficients |
| --- | --- | --- | --- | --- | --- |
|  |  |  |  |  | microbial relative abundance |
| Maternal age | 1.182 | 0.347 | 0.0372 | 0.56 | *Bacteroides* (0.69), *Bifidobacterium* (-0.45), *Clostridium_sensu_stricto* (1.11), *Flavonifractor* (0.64), *Veillonella* (0.50), *[Ruminococcus]_gnavus_group* (1.31) |
| Milk components: |  |  |  |  |  |
| Leptin | 1.518 | 0.230 | 0.0450 | -0.05 | *Bacteroides* (-0.06), *Veillonella* (0.04) |
| Cortisol | 1.511 | 0.232 | 0.0444 | 0.62 | *Bacteroides* (-1.23), *Bifidobacterium* (-0.79) |
| IgA | 1.459 | 0.246 | 0.0401 | 12.1 | *Clostridium_sensu_stricto_1* (-29.9), *[Ruminococcus]_gnavus_group* (6.49) |
| IgM | 1.398 | 0.255 | 0.0568 | -0.64 | *Bacteroides* (-3.09), *Bifidobacterium* (2.64), *Escherichia-Shigella* (2.11), *Veillonella* (0.23). |

The table shows the results of five multivariable linear regression models performed using data from bacterial genus with $RA \geq 1\%$ in the stool samples (n=34). The type of birth was included as a covariate in all models. Variables considered in the models were those that showed $r > |0.2|$ in the Spearman correlation analysis of the whole cohort. Values within brackets indicate the beta coefficient for each variable.

## DISCUSSION

Our results show that the gut microbiota composition of exclusively breastfed infants at an average 5.4 months of age still exhibits differences in the relative abundance of taxa (up to the genus level) depending on whether they were delivered vaginally or by cesarean section. Studies conducted in different regions of the world have reported variable persistence of microbiota differences depending on the mode of delivery^53^. Importantly, unlike other reported cohorts, all infants in our study were full-term and exclusively breastfed at the time of sampling, which may contribute to the different outcomes reported. In contrast to Cabrera-Rubio et al. (2012)^54^, we did not find significant differences depending on whether the cesarean section was elective or preceded by labor; however, this effect cannot be completely ruled out given the small number of samples in each subgroup.

With respect to specific microbial profiles, higher Firmicutes-Bacteroidetes ratios have been associated with excessive weight gain and obesity ^55–57^. Moreover, the families *Enterobacteriaceae* and *Clostridiaceae* have been associated with a higher risk of gut inflammation and adiposity^58,59^. Based on this background, our results on microbiota composition may suggest an influence of delivery mode on metabolic trajectories and the risk of obesity at later stages. However, this study does not include the complementary feeding period or the changes in microbiota composition at older ages. During our study period we did not observe significant differences in weight gain between the two groups, the proportion of infants with weight-for-age percentiles above 85 was 30% and 21% for the V and CS groups, respectively. A longer follow-up, including more specific parameters of infant growth and metabolism, could yield interesting data.

The variation in breastmilk composition could be a factor contributing to the acquisition of shifts in bacterial composition from mothers to infants ^9^. Fatty acids and human milk oligosaccharides (HMOs) profiles are particularly important components in defining the predominant bacterial types and the metabolites they generate, which exert diverse local and systemic effects^60,61^. Although we did not find significant differences in the macronutrient content of milk between the two groups, it is noteworthy that different associations with taxa at the family and genus levels were observed in groups CS and V. While both groups showed a positive correlation between fat content and families or genera belonging to the phylum *Proteobacteria*, in group CS the relative abundance of members of the phylum *Firmicutes* appeared to show an opposite trend.

Maternal diet may indirectly influence the composition of infant microbiota, particularly because fat is the milk macronutrient most dependent on maternal intake^62^. All mothers reported an unrestricted omnivorous diet during pregnancy and the postpartum period. A limitation of this study is the lack of a detailed record of maternal dietary intake, which could contribute to the results obtained.

Our results regarding the RA of *Bifidobacterium* in both groups differ from other studies reporting a higher predominance of these bacteria at both the family and genus levels in the fecal microbiota of groups of infants including both exclusively and mixed breastfeeding ^6,15,63^. In contrast, our study, which included only exclusively breastfed infants, shows results comparable to those obtained in other cohorts with similar characteristics in relation to the type of breastfeeding^61,64^.Other studies have reported a higher abundance of certain *Bifidobacterium* species in the fecal microbiota of infants born vaginally compared with those delivered by cesarean section ^65^. Although we did not find differences in *Bifidobacterium* abundance at the family or genus levels according to delivery mode, the association observed with total carbohydrate content exclusively in group CS is noteworthy. These results encourage further investigation into the characterization of specific types of HMOs in the milk of Uruguayan women, because the secretory status was shown to impact the microbiome profile in CS-delivered infants ^66,67^.

The maternal immune system, and particularly the IgA in the infant’s intestine, contributes to shaping the microbiota composition and the immune system. In addition to pathogen clearance, IgA promotes the colonization of beneficial microbes, contributes to microbial diversity and acts as a nutrient source for certain taxa ^68^. At six-month age, all the IgA in the infant gut derives from breast milk, and is mostly associated with bacteria, with the genus *Bifidobacterium* being one of the main binding targets ^69^. In our study, we did not find differences in total IgA content in milk or in the relative abundance of *Bifidobacterium* between groups V and CS. However, the positive correlation between IgA and *Bifidobacterium* observed only in group V suggests that these bacteria may be differently coated with IgA.

In parallel with this difference, we found that only in group V there was a positive correlation between milk IgA levels and the relative abundance of *Bifidobacterium* at both the family and genus levels, with a similar trend observed in fecal samples. Considering these observations together, it cannot be ruled out that maternal physiological differences associated with the peripartum period, such as variations in bacterial translocation, which may affect the initial bacterial inoculum in milk, and the migration of antibody-producing cells to the mammary gland^70^ could influence microbiota homeostasis and its fine composition. Furthermore, the possible effect of antibiotic administration during cesarean deliveries cannot be excluded.

The correlation observed between the relative abundance of *Flavonifractor* and maternal perceived stress in group CS also deserves further investigation, as previous studies have reported an association between the abundance of this genus and psychological conditions involving stress ^71^.

A major strength of our analysis is the use of a homogeneous cohort, which minimizes the influence of factors that could strongly affect the infant microbiota, such as parity, gestational age, and type of breastfeeding. Nevertheless, the limited sample size could contribute to the lack of significant findings in the multivariable models.

The higher maternal age observed in the CS group aligns with national data indicating increased cesarean delivery rates with advancing maternal age. Nevertheless, in the 30–34-year age range (n =20), the percentage of cesarean (20%) was lower than the national average ^72^. The recruitment strategy may have contributed to the inclusion of participants with a high sociocultural level (88% with tertiary education); therefore, the present findings should not be generalized to the general population.

In summary, this first report on Uruguayan infants during exclusive breastfeeding shows that gut microbiota composition remains influenced by delivery mode and may be differentially modulated by breast milk bioactive and/or nutritional components and maternal perceived stress. Our results contribute to a better understanding of how maternal and perinatal factors influence the development of the early microbiota. A larger sample size and an extension to groups with diverse maternal sociocultural backgrounds are among our future perspectives.

## Acknowledgments

We are deeply grateful to the mothers participating in this work for their invaluable collaboration. We thank Karina Machado, MD, for her guidance in the description of the infant growth parameters. CM, JRSS, CRC, and AH are researchers of the Sistema Nacional de Investigadores (SNI) of the Agencia Nacional de Investigación e Innovación (ANII, Uruguay).

## Funding

This work was supported by the Comisión Sectorial de Investigación Científica – Universidad de la República under Grant CSIC (301/2020 to AH). Additional support was provided by Programa de Desarrollo de las Ciencias Básicas (PEDECIBA), Uruguay to CM, JRSS, CRC, and AH.

## Author contributions

Conception and design: AH, JRSS; acquisition of data: JP, LHA, CM, JPP, LB, AP, CRC; analysis and interpretation of data: AH, JRSS, JPP, CF, CRC, CM, LB, LHA; drafting the article or revising it critically for important intellectual content: AH, JRSS, CM, LHA, JPP. The final approval of the version to be published: all authors.

## Competing interest

The authors declare no conflict of interest.

## Consent statement

All patients provided informed consent to participate in this study, which was approved by the Ethics Committee of Facultad de Química, Universidad de la República, (Exp. No 101900-000659-20) in accordance with the Declaration of Helsinki.

## Notes

### Competing Interest Statement

The authors have declared no competing interest.

### Summary of Updates

Author order updated since it was wrong, CRC goes before JRSS

