## Supplementary Figure 1 and Supplementary Table 1 for "Links Between Gut Microbiota of Uruguayan Infants, Breastmilk Composition, and Maternal Factors During Exclusive Breastfeeding"

Supplementary Figure 1. Rarefaction curves of ASV richness by delivery mode.

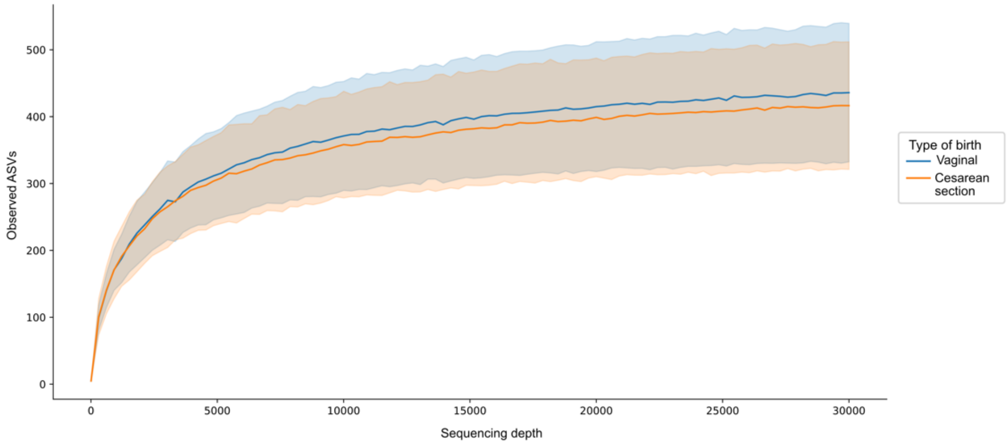


Rarefaction curves depict the relationship between sequencing depth and observed amplicon sequence variant (ASV) richness in stool samples from infants born by vaginal delivery or cesarean section. Solid lines represent the mean number of observed ASVs for each delivery mode, Shaded areas correspond to the standard deviation, indicating within-group variability. Across the full range of sequencing depths evaluated, samples from vaginally delivered infants consistently show higher mean ASV richness than those from cesarean-delivered infants.

**Supplementary Table 1. Simple correlation analysis.**

| Variables | | | |
|---|---|---|---|
| *Mothers* | *RA (%)* | r | P-value |
| Age | *g__Bacteroides* | -0.3226 | 0.059 |
| | *g__Bifidobacterium* | -0.2160 | 0.213 |
| | *g__Clostridium_sensu_stricto_1* | 0.2459 | 0.154 |
| | *g__Flavonifractor* | 0.2273 | 0.189 |
| | *g__Veillonella* | 0.2393 | 0.166 |
| | *g__[Ruminococcus]_gnavus_group* | 0.3770 | 0.026 |
| Perceived stress | *g__Veillonella* | 0.2056 | 0.236 |
| *Milk components* | | | |
| Cortisol | *g__Bacteroides* | -0.2331 | 0.178 |
| | *g__Bifidobacterium* | -0.2690 | 0.118 |
| IgA | *g__Clostridium_sensu_stricto_1* | -0.2938 | 0.087 |
| | *g__[Ruminococcus]_gnavus_group* | 0.2510 | 0.146 |
| IgM | *g__Bacteroides* | -0.3206 | 0.060 |
| | *g__Bifidobacterium* | 0.2742 | 0.111 |
| | *g__Escherichia-Shigella* | 0.2156 | 0.213 |
| | *g__Veillonella* | 0.2050 | 0.238 |
| Leptin | *g__Bacteroides* | -0.2196 | 0.205 |
| | *g__Veillonella* | 0.3946 | 0.019 |
| Adiponeptin | *g__Flavonifractor* | -0.3113 | 0.069 |
| *Infants* | | | |
| IgA in stool | *g__Bifidobacterium* | 0.3003 | 0.080 |

The table shows the results of Spearman correlation analyses between the relative abundance (RA) of bacterial genera and maternal variables, as well as biochemical parameters measured in stool and milk samples. Only genera with RA ≥ 1% were included in the analysis, and correlation coefficients |r| > 0.2 are shown for the entire cohort (n = 34).
